# Representational geometry of punishment learning across basolateral amygdala neurons

**DOI:** 10.64898/2026.10.25.761607

**Authors:** Philip Jean-Richard-dit-Bressel, Nikki Huang, Eun A Choi, E. Zayra Millan, Cassandra V. Occelli Hanbury-Brown, A. Simon Killcross, Gavan P. McNally

**Author notes:** Corresponding author. Abbreviated title: Punishment learning.

## Abstract

Punishment learning allows animals to suppress actions that cause harm while maintaining actions that earn reward. Although the basolateral amygdala is necessary for punishment learning, how its population activity changes across learning, and how this activity reflects an action’s consequences, remain unknown. We used longitudinal calcium imaging to track 1,147 basolateral amygdala neurons in male mice across instrumental training, punishment, choice test, and punishment extinction. Punishment learning progressively transformed basolateral amygdala neuron activity associated with the punished action. This activity diverged from that associated with the unpunished action and became aligned with activity evoked by the punisher itself. This alignment was action and contingency specific. It preserved the direction of individual neurons’ responses to the punisher: neurons activated by the punisher become more activated by the punished action alone, whereas neurons inhibited by the punisher become more inhibited by the punished action alone. We then asked whether reward and punishment were encoded separately or collapsed onto a common value dimension across basolateral amygdala neurons. Punishment added a strong shock component to action-related activity without reducing its reward component, while reward and shock themselves occupied largely orthogonal population directions. Thus, during punishment learning, basolateral amygdala population activity can encode opposing consequences of the same action instead of simply changing what that action is worth.

**Significance statement:** Learning to avoid actions that cause harm is essential for adaptive behavior, but how the brain represents the changed value of a punished action is poorly understood. By tracking the same basolateral amygdala neurons across punishment learning, choice, and extinction, we show that the representation of an action progressively incorporates the representation of its aversive consequence. This transformation is specific to the punished action and reverses when the action–punisher relation is extinguished. Importantly, punishment does not simply move the action along a single value dimension. Instead, reward and shock occupy separable dimensions of amygdala population activity. These findings reveal a representational architecture in which punishment changes what an action is expected to cause instead of just changing what it is worth.

---

Punishment learning is essential for adaptive decision-making. When an action produces aversive consequences, animals must use this information to suppress that action while maintaining other behaviors that earn reward (Jean-Richard-Dit-Bressel et al., 2018; Piantadosi et al., 2021). Much of what is known about the neural representation of aversive learning comes from Pavlovian fear conditioning, in which external cues predict aversive events and elicit involuntary defensive responses (McNally et al., 2011; Herry and Johansen, 2014). Longitudinal imaging of basolateral amygdala (BLA) populations has shown that fear conditioning transforms the representation of a conditioned stimulus toward that of the aversive unconditioned stimulus, and that this relationship changes again during extinction (Grewe et al., 2017; Zhang and Li, 2018). These findings show how an aversive outcome can transform the BLA representation of a stimulus that predicts it. Whether an animal’s own voluntary actions undergo a comparable transformation when they cause the aversive outcome remains unclear.

In punishment learning, animals must detect the relation between an action and an aversive outcome, integrate that outcome with the reward earned by the same action, and selectively suppress the punished action while maintaining other rewarded behaviors (Bolles et al., 1975, 1980; Varas et al., 2026). Lesion, reversible inactivation, and optogenetic silencing experiments show that BLA is necessary for this learning and its expression (Killcross et al., 1997; Jean-Richard-Dit-Bressel and McNally, 2015; Orsini et al., 2015; Truckenbrod et al., 2023; Gregory et al., 2026; Piantadosi et al., 2026). Moreover, fibre photometry of calcium transients shows that BLA population activity tracks aversive outcomes and the actions that cause them (Jean-Richard-dit-Bressel et al., 2022), while recent calcium imaging shows rapid reorganization of BLA choice representations as previously established punishment risk changes within a session (Piantadosi et al., 2026). What remains unknown is how the representation of a specific action evolves as its relation to punishment is learned, expressed, and extinguished.

Two contrasting accounts suggest different ways in which punishment could alter an action’s representation. In scalar reinforcement-learning, appetitive and aversive outcomes contribute with opposite signs to a common action value that is used to guide action selection (Sutton and Barto, 2018). If BLA population activity represents this integrated value, punishment learning might shift the action representation along a shared appetitive-to-aversive dimension. This possibility is consistent with evidence that primate and rodent BLA neurons encode positive and negative value (Paton et al., 2006; Belova et al., 2008; Namburi et al., 2015; Kim et al., 2016; Tye, 2018). Alternatively, BLA could represent the specific outcomes of an action, allowing punishment to evoke representations of its aversive consequence without altering representations of rewarding outcomes. This possibility accords with the proposal that BLA represents the incentive properties of specific motivationally significant outcomes rather than just their general affective value (Balleine and Killcross, 2006).

Here we used longitudinal calcium imaging to track the same BLA neurons across weeks as mice progressed through instrumental training, punishment, choice between punished and unpunished actions, and punishment extinction. We asked what kinds of task-related activity patterns occur across BLA neurons across these phases and how this related to behavior. We also asked whether learning selectively reorganizes the representation of the punished action, whether this reorganization persists in the absence of punishment, and whether it reverses during extinction. Using BLA population responses to shock and reward as references, we then tested whether the punished action shifts along a common appetitive-to-aversive dimension or acquires a separable representation of the punisher while retaining its representation of reward.

## Materials and Methods

### Subjects

Male C57BL/6J mice (Australian Resources Centre; 8 - 10 weeks old) were housed in ventilation racks on corn cob bedding in a climate-controlled colony room under a 12:12 hr light/dark cycle. Female mice were not available from the supplier during this experiment. Mice had free access to food and water until 2 days prior to commencement of behavioral training. For the remainder of the experiment, they received 1 - 2 g of food and free access to water each day. This experiment was approved by the UNSW Animal Care and Ethics Committee and performed in accordance with the Animal Research Act 1985 (NSW), under both ARRIVE guidelines and the National Health and Medical Research Council Code for the Care and Use of Animals for Scientific Purposes in Australia (2013).

### Surgery

Mice were anaesthetized using 5% isoflurane in oxygen-enriched air, then fixed into a stereotax (Model 1900, Kopf Instruments). Mice were maintained on 2.5% isoflurane during surgery. 0.1 ml Marcaine (0.5%) was injected subcutaneously at the incision site. Ophthalmic gel (Viscotears, Alcon) was applied to avoid eye-drying. Mice received subcutaneous injections of antibiotic (Duplocillin, 0.15 ml/kg of body weight subcutaneously) and 5 mg/kg carprofen (Rimadyl, Zoetis) immediately post-surgery.

0.5μl AAV9-hSyn-GCaMP7f (pGP-AAV-syn-jGCaMP7f-WPRE was a gift from Douglas Kim & GENIE Project (Addgene viral prep # 104488-AAV; http://n2t.net/addgene:104488; RRID:Addgene 104488) and was unilaterally injected using a 30-gauge conical-tipped microinfusion syringe (SGE Analytical Science) connected to a UMP3 with SYS4 Micro-controller microinjection system (World Precision Instruments). BLA coordinates were 1.5 mm posterior, ± 3.4 mm lateral and 4.8 mm ventral from bregma (Paxinos and Franklin, 2019). The GRIN lens (0.5 mm diameter) was implanted 0.5 mm above. The lens was integrated into a baseplate that was secured to the skull using superglue and dental cement. Mice were allowed 6 weeks post-surgery recovery during which their condition was monitored daily (first week) then three times weekly (for the remainder of the experiment). Mice were single-housed post-surgery to prevent damage to GRIN lens implants. From 3 - 6 weeks post-surgery, mice were connected to the Miniscope weekly for habituation.

### Punishment task and calcium imaging

All behavioral training and testing occurred in ENV-307A Standard Modular Chambers for Mouse (Med Associates). Each chamber was fitted with two retractable levers located either side of magazine for delivery of 20 mg grain pellets (BioServ, Dustless Precision Pellets). The chambers had a stainless-steel grid floor through which the punisher was delivered and were located in sound attenuating cabinets.

6 – 7 weeks after surgery, mice underwent a punishment task (**Figure 1B**). Mice were initially trained to press two levers (Action 1 [A1] vs Action 2 [A2]) for food. Mice first received 2 sessions of fixed-ratio 2 (FR2) lever-press training. Both levers were extended, and every second press on either lever was reinforced with delivery of a pellet to an external magazine. Each lever retracted after being pressed 25 times. The session ended after 50 min or when both levers had retracted. Mice then received daily 40-min lever-press training sessions over the next 10 days. Each lever was presented individually in 5 min alternating blocks; there were four blocks per lever across 40 min. Lever-pressing was reinforced with pellet delivery on a 30-s variable-interval (VI30s) schedule. The first lever to be extended (left or right) was fully randomised.

**Figure 1.**
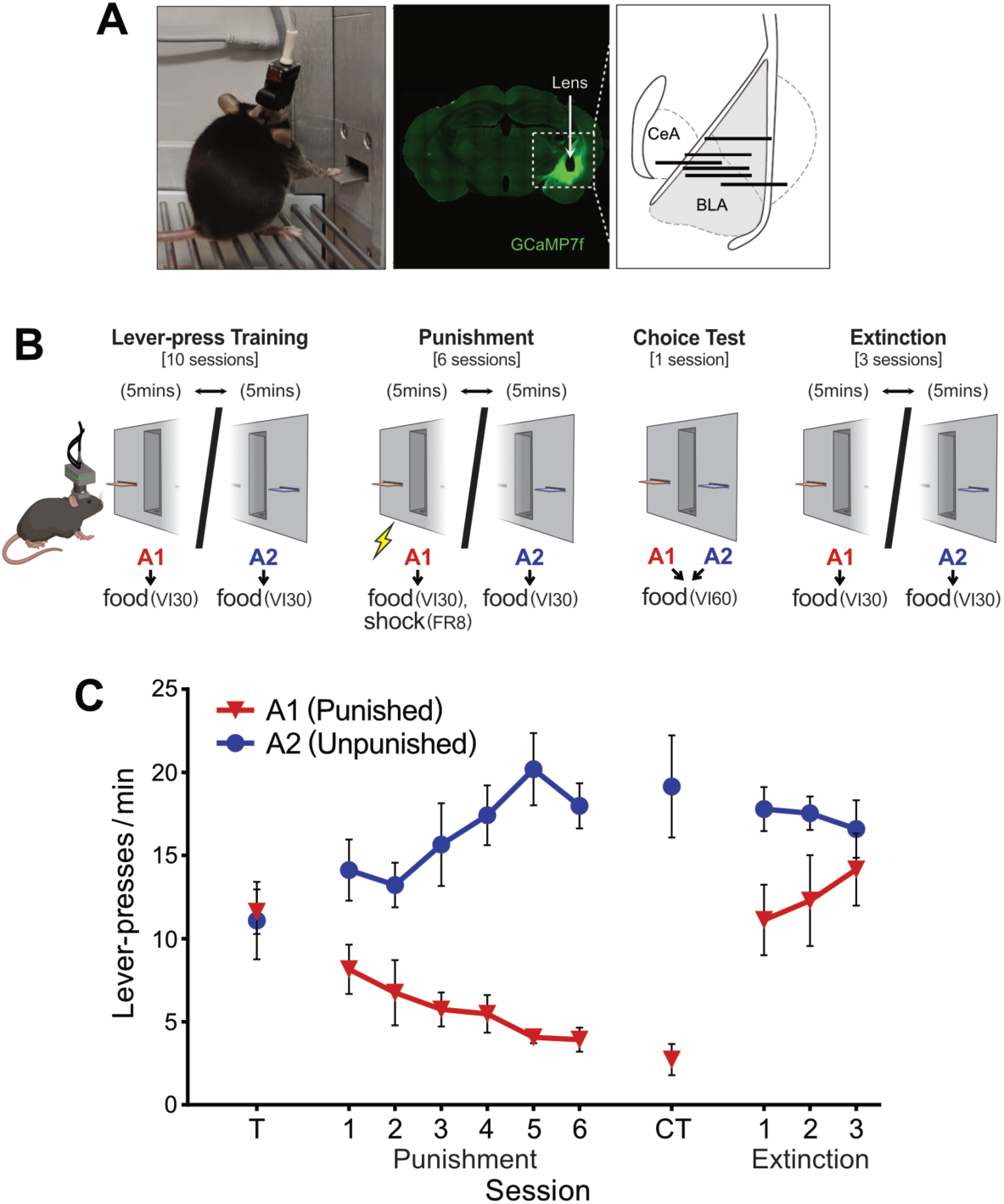
Longitudinal imaging of BLA neurons across a punishment task. **A** Left: Photo of miniscope-tethered mouse during punishment task. Middle: Coronal brain slice showing the GCaMP7f expression and GRIN lens track in BLA. Right: Schematic of coronal mouse brain section with reconstructed positions of the GRIN lens implant (black lines; n = 6) **B**, Punishment task phases. Mice learned to press two individually-presented levers (A1,A2) for food pellets during Training. A shock-punisher contingency was introduced to A1 in Punishment sessions. Mice were then given an unpunished choice test where both levers were concurrently available. Mice then received punishment extinction sessions (identical to pre-punishment training). **C**, Mice showed expected changes in behavior. Prior to punishment (T), mice had no preference between levers. Punishment drove a selective suppression of punished A1 responding. A strong preference for A2 over A1 was observed in choice test (CT). Avoidance of A1 was gradually lost across extinction sessions.

Next, mice received punishment phase sessions. Punishment sessions were identical to VI30s lever-press training sessions, except presses on the designated punished lever (A1) delivered a 0.5 s shock on a FR8 schedule independent of pellet outcome. Presses on the other lever (A2) remained unpunished. To preserve opportunities to measure shock- and A1-related neural activity, shock intensity was gradually increased across punishment training (Days 1-2: 0.2 mA; Days 3-4: 0.3 mA; Days 5-7: 0.4 mA).

Following punishment phase, there was a single 10-min choice session that involved concurrent presentations of both levers. During this choice session, lever presses were reinforced with pellet delivery on a unified VI60s schedule and there was no punishment. Finally, mice received 3 x 40-min punishment extinction sessions that were identical to initial VI30s training as described above (shocks no longer delivered).

Mice were connected to the miniscope daily throughout the experiment except on Days 3 - 5 of VI30s lever-press training, when mice were untethered to accelerate lever press acquisition. Recordings were made for key segments of each experiment session. On VI30s lever-press training, all punishment training days, and all punishment extinction days, recordings were made for the first and last 10 min of the session. Recordings were made for the entire 10 min of the choice test. Fluorescence Ca^2+^ imaging videos were acquired using nVista HD software (Inscopix Inc, Mountain View, CA, USA) with a field of view of approximately 600 x 600 μm, LED power of 0.3-0.5 mW/mm^2^, analogue gain of 1-3, and 2x spatial down sampling. 12-bit images (1,000 x 1,000 pixels) were acquired at a frame rate of 30 Hz, with each pixel covering 0.6 x 0.6 mm in tissue. For each mouse, the same imaging parameters were used for all imaging sessions. The video data were streamed directly to a hard disk (90 -100 MB/s).

### Histology

At the end of the experiment, mice were deeply anaesthetized via IP sodium pentobarbital (100 mg/kg) and perfused with 0.9% saline solution containing 1% sodium nitrite and heparin (5000 IU/ml), followed by phosphate buffer solution (0.1 M) with 4% paraformaldehyde. Brains were extracted, sliced coronally (40 μm) using a cryostat and processed for immunofluorescence to identify location of GRIN lens and GCaMP expression.

### Experimental design and statistical analysis

#### Behavior

The primary behavioral variable of interest was lever presses per minute. These data were analyzed using repeated measures ANOVA.

#### Calcium imaging preprocessing

For each mouse, recordings from the same day were spliced into a single file and motion-corrected. Each image frame of the motion-corrected video was re-expressed in units of relative change in fluorescence, Δ*F* = (*F*(*t*) – *F*0) /*F*0, where *F*0 is the mean brightness per pixel across a recording. Individual candidate BLA neurons were then identified via convex non-negative matrix factorization with spatiotemporal constraints (CNMFE).

CNMFE parameters of cell diameter, minimum pixel correlation, and minimum peak-to-noise ratio were manually determined for each mouse to prevent over- or under-counting of cells. All other CNMFE parameters were set to default values. CNMFE-derived Δ*F*/*F* was low-pass (3 Hz) filtered to remove high-frequency noise identified via Fast Fourier Transform.

For each mouse, a longitudinal registration algorithm within Inscopix Data Processing Software was used to longitudinally register cell across sessions. There were 1,147 longitudinally registered cells across 6 mice (Mouse 1–6 = 139, 235, 196, 203, 111, 263 cells, respectively). Average number of sessions tracked per cell was 3.12 sessions (subject range = 2.11 – 4.05). Late lever-press training sessions (T9 – 10; “T”) and late punishment sessions (“Pun 4-6”) were concatenated to pool more data from these behaviorally stable phases.

Cell activity and longitudinal registration data were exported for further analysis in MATLAB. For each cell, low-pass filtered Δ*F*/*F* from -5 to +7 s around events of interest were collated. Events were A1 alone (i.e., presses on punished lever not yielding shock or pellet), A2 alone (i.e., presses on unpunished lever not yielding pellet), shock alone (i.e., those that did not co-occur with pellet delivery, although necessarily co-occurring with A1) and reward (i.e., pellet consumption). The -5 to 7s window was chosen to capture a sufficiently broad window of peri-event activity. Each event trial was re-baselined to pre-event activity (−5 to -3 s) and normalized by its sum squared deviation from zero (nullZ) (Keevers and Jean-Richard-dit-Bressel, 2025) to minimize the influence of changing baseline and outlier trials. Peri-event activity per cell was calculated as mean normalized activity across trials; these averaged peri-event cell responses were used in all further analyses.

#### Clustering and event-related transient analyses

To identify neuronal ensembles based on cross-task activity, K-means clustering was performed on peri-event activity of cells co-registered across late lever-press training, early punishment (Pun 1), and late punishment (Pun4-6) (*n* = 154 cells). K = 2 – 8 cluster solutions were examined. A 3-cluster solution was confirmed as optimal; all cells in this solution had positive silhouette values, indicating each cell’s activity profile was well-represented by their cluster. Clusters were differentiated by activity patterns to shock, reward, and punished action (A1) during late punishment. Additional cells (those only partially co-registered across sessions) were incorporated into clusters if they had positive silhouette values for these transients across shared sessions. Following this, Cluster 1 comprised 165 cells, Cluster 2 comprised 270 cells, and Cluster 3 comprised 176 cells. Significance of peri-event activity per cluster was identified via 95% bootstrapped confidence intervals (Jean-Richard-dit-Bressel et al., 2020) to identify when activity across cells in a cluster significantly deviated from pre-event baseline, and when activity significantly differed between specific events (within-subject difference waveform for: shock vs reward, A1 vs A2).

#### Cross-session similarity analyses per ensemble

Within each cluster, similarity between A1, A2, shock, and reward activity kernels was quantified by deriving a normalized fit score for each pairwise kernel comparison. The normalized fit score (−1 to 1) was calculated as the dot product of two peri-event kernel transient vectors (each normalized according to its sum square deviation from zero). A positive fit score indicates that two kernel transients deviate from baseline in a similar way across the peri-event interval. A negative fit score indicates that two kernel transients deviate from baseline in inverse ways. Zero indicates that two kernel transient vectors are orthogonal to one another.

To determine whether peri-event kernel transients across sessions were significantly similar (fit > 0) or inverse (fit < 0) to each other, 95% CI limits for fits were obtained from 2.5 and 97.5 percentiles of bootstrapped fit distributions (fits of randomly resampled mean transients, 1000 iterations). Two transients were deemed significantly similar if the fit CI was entirely above zero, or significantly inverse if the fit CI was entirely below zero. To visualize the overall geometry of similarity/dissimilarity between peri-event activity kernels across sessions, fit scores were converted into fit distances (1 - fit score) and arranged in 2D space via multidimensional scaling (MATLAB *mdscale* function, criterion = metrics-stress).

#### Relationships between ensemble activity and behavior

To examine relationships between action recoding per ensemble and behavior change across phases, we computed fit scores (as described above) for an action’s kernel for a session against late punishment action versus pre-punishment action kernels (Relative fit score = Pun4-6 fit – Training fit). Positive values indicate that an action’s activity pattern more closely resembles late-punishment than pre-punishment patterns, whereas negative values indicate an activity pattern is more similar to training than late punishment. This was computed per subject per cluster. In this way, each mouse had self-normalized relative fit scores per cluster for each action and session (excluding sessions used to anchor fits [Training, Pun 4-6]).

Suppression ratios provided a self-normalized index of behavior change per lever by quantifying change in lever-press rates relative to pre-punishment. Suppression ratios were calculated per lever and session as: LP rate / (LP rate + Training LP rate) per lever. Values range from 0 to 1, where 0.5 indicates no change relative to late lever-press training, values >0.5 indicate higher punished-lever-press rates, and values <0.5 indicate lower rates. Relationships between relative fit scores and suppression ratios per lever were assessed using linear regression.

#### Whole-population geometry

Each neuron’s response to each event was summarized as a single scalar: the mean normalized peri-event transient over the 0–1 s window following event onset, averaged across trials of that type. This yielded one value per neuron per event-by-session condition, giving a matrix of 20 conditions by 1,147 registered cells.

Each condition was treated as a vector in neuron space. Population responses were described by two quantities: magnitude, defined as the root-mean-square response across neurons, and direction, compared between conditions using cosine similarity. Because cosine similarity is invariant to vector length, these measures distinguish changes in the overall strength of the population response from changes in its pattern across neurons. Not every neuron could be registered across every session: only 12 of the 1,147 neurons were registered in all six sessions. Analyses therefore used the neurons available for the sessions being compared. Within-session comparisons used exactly the same neuron set. Cross-session comparisons used neurons registered in both sessions, giving 63–269 neurons per pair. An additional control was required when comparing cosines across sessions because the two cosines could contain different neurons. For example, shock alignment during lever-press training was calculated from 265 neurons and shock alignment during Punishment 4–6 from 564 neurons, but only 154 contributed to both estimates. We therefore repeated each such comparison using only neurons contributing to both cosines, ensuring that any difference in cosine similarity was confirmed in the same populations (n = 88–564).

All resampling used a fixed seed. Confidence intervals were 95% percentile bootstraps over neurons with 5,000 resamples. The significance of a single cosine similarity was assessed by permuting neuron identity in one vector, destroying neuron-by-neuron correspondence while preserving both marginal distributions (5,000 permutations). Differences between cosine similarities were bootstrapped over the neurons contributing to both cosines, with the two cosines resampled together (5,000 resamples; 10,000 for correlation contrasts). P values were computed as (k + 1)/(B + 1) after North et al. (2002), where k is the number of resamples at least as extreme as the observed value and B is the number of resamples. This bounds p at 1/(B + 1), approximately 2 × 10^−4^; values at this bound are reported as p < 0.0002. The Benjamini–Hochberg false-discovery-rate correction was applied across the entire family (Benjamini and Hochberg, 1995). For visualization only, the 20 conditions were embedded by multidimensional scaling of cosine distances computed using pairwise-complete neurons; the first two dimensions captured 46% of the absolute eigenvalue mass.

Alignment with the punisher was quantified as each condition’s cosine similarity with a fixed shock vector, computed separately against shock vectors from Pun 1 and Pun 4–6. For additional shock-tuning analyses, neurons were ranked by their Pun 4-6 shock response and A1 response magnitude was averaged per shock response quintile. Three controls tested alternative explanations for the resulting gradient: correlation of absolute values tested a gain or excitability account; partial correlation controlling for reward response tested a salience account; and Spearman correlation and correlation after trimming the most extreme 20% of shock responders.

The learning-induced change in the punished-press vector was calculated as the Pun 4–6 vector minus the lever-press training vector, using neurons imaged in both sessions, and this displacement was projected separately onto shock and reward vectors. A single value axis predicts that a displacement with a positive shock component should have an oppositely signed reward component of equal magnitude. The angular separation of the shock and reward vectors was additionally expressed after correction for axis reliability, estimated from cross-session self-similarity (shock, 0.48; reward, 0.39).

## Results

### Histology

GCaMP7f was expressed in BLA and GRIN lenses had optical focal planes primarily within the BLA for all mice (**Figure 1A**).

### Behavior

Mice rapidly learned punishment (**Figure 1B**). Across six days of punishment training, mice learned to selectively suppress the punished action (Lever: F(1,5) = 56.300, p = .001; Session: F(1,5) = 0.385, p = .562); Lever*Session: F(1,5) = 54.585, p = .001). During choice test, when A1 and A2 were concurrently available in the absence of punishment, mice strongly preferred the safe lever over the previously punished lever (F (1, 5) = 25.054, p = 0.002). Subsequently, across three days of extinction training in which the shock punisher was omitted, mice flexibly updated their learning and selectively increased responding on A1 (Lever: F(1,5) = 8.505, p = .033; Session linear trend: F(1,5) = 0.302, p = .606; Lever*session: F(1,5) = 6.514, p = .051).

### BLA imaging

We longitudinally imaged calcium activity in BLA neurons of six mice across the task. In total, 1,147 unique neurons were registered across recording sessions (192 +/-23 neurons per mouse). Individual neurons were not always observed in every session, so the number of cells available for analysis varied by session: 508 were present during late lever-press training (T), 315 during Pun 1, 573 during Pun 4–6, 267 during the choice test, 291 during Ext 1 and 269 during Ext 3.

### Profiles of BLA task-related activity

*k-*means clustering was applied to peri-event activity around actions (A1, A2) and outcomes (reward, shock) across late lever-press training, early punishment (Pun 1) and late punishment (Pun 4–6). Clusters were thus defined by their joint responses to both actions and outcomes across sessions. Clusters were initially identified using the 154 neurons registered across all three phases. Silhouette analysis across K = 2–8 favored a 3-cluster solution. In this solution, every neuron had a positive silhouette value, indicating that each neuron had strong within-cluster cohesion and between-cluster segregation. Neurons not registered across all three stages were then assigned to the nearest of these three profiles, giving 611 of 1,147 neurons a cluster identity: Cluster 1 (*n* = 165), Cluster 2 (*n* = 270) and Cluster 3 (*n* = 176).

The three clusters differed profoundly in their responses to reward and punishment. Cluster 1 neurons showed a biphasic excitatory-then-inhibitory response to reward and robust inhibition to shock (**Figure 2A**). Cluster 2 neurons showed the opposite: robust excitation to shock and inhibition during reward consumption (**Figure 2B).** Cluster 3 neurons were strongly excited by reward and weakly excited by shock (**Figure 2C**). This opposing organization is consistent with previous reports of valence-selective populations in BLA (Namburi et al., 2015; Beyeler et al., 2016; Kim et al., 2016; Piantadosi et al., 2024) Interestingly, reward responses in Clusters 1 and 3 were attenuated during Pun 1 relative to all other stages, suggesting that the initial punishment is a distinct phase in which reward-related BLA activity is transiently altered.

**Figure 2.**
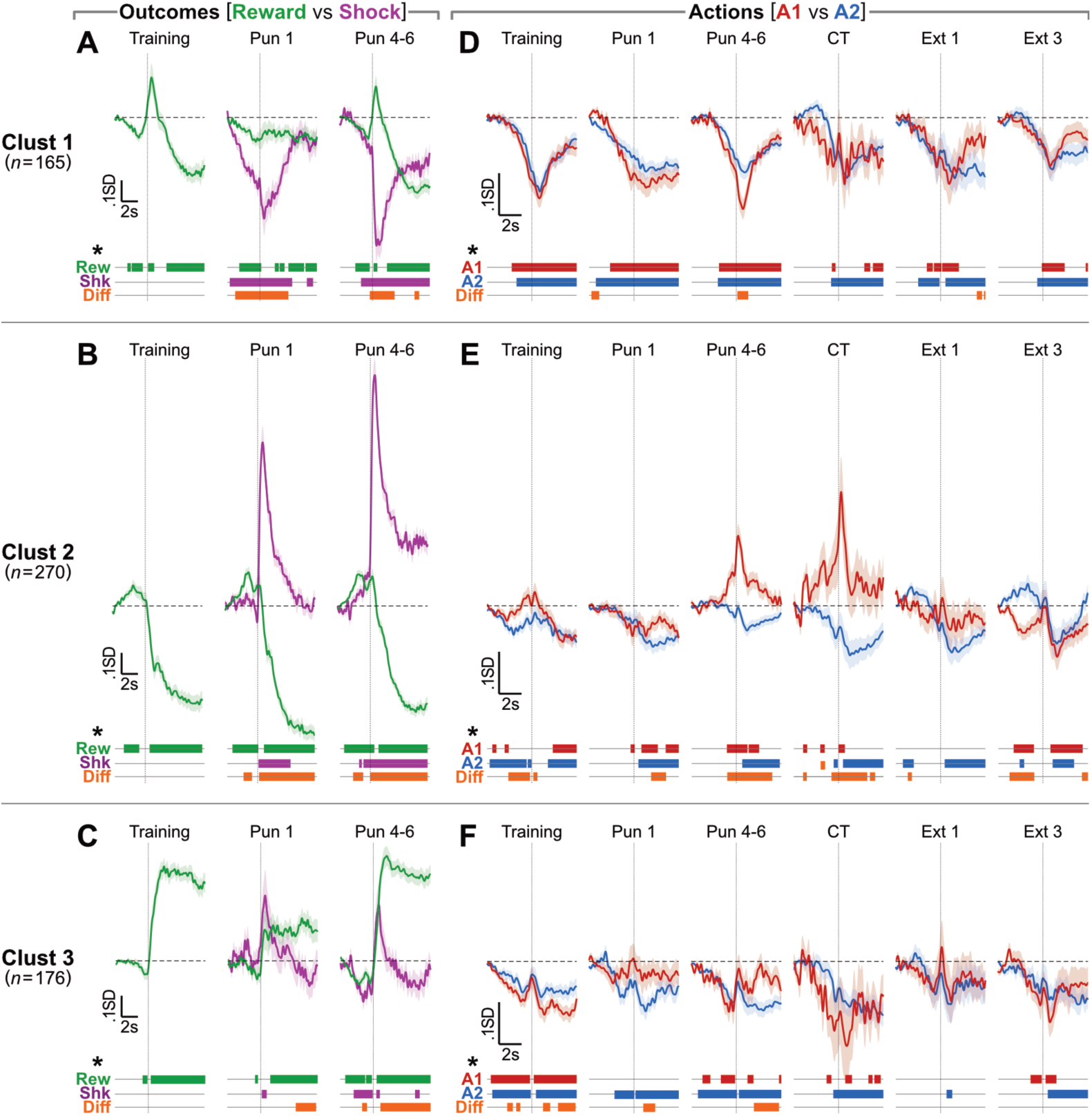
Key profiles of BLA neuron activity to outcomes and actions. A-F Mean ± SEM normalised activity transients around outcomes (reward consumption [green] vs shock [purple]; A-C) and actions (A1 [red] vs A2 [blue]; D-F) across k-means-identified clusters (Clust 1-3). Horizontal dashed lines per panel indicate dF/F = 0 (pre-event baseline). Vertical dotted line per panel indicates event onset. Colored bars beneath each panel indicate when peri-event transients significantly differed from 0 (baseline) or between plotted events (Diff), as determined via 95% bootstrapped confidence intervals.

We next examined activity around the two actions (using trials without outcome delivery). Across clusters, actions were generally associated with inhibitory transients (**Figure 2D–F**). Cluster 2 was the exception (**Figure 2E**). During lever-press training and early punishment, Cluster 2 neurons were inhibited around both punished A1 and unpunished A2 presses. However, by late punishment they showed phasic excitation around A1 presses while A2 presses continued to evoke inhibitory transients. This dissociation persisted into the choice test, where no shocks were delivered, indicating that it reflected learning about the punished action rather than within-session shock delivery. During extinction, Cluster 2 activity around A1 presses reverted to inhibitory transients.

### Representational geometry within BLA ensembles and its relation to behaviour

We next examined coding principles of each ensemble by applying similarity analysis to each cluster’s activity kernels for actions and outcomes across sessions. Similarity of activity between pairs of task events by session was calculated to produce a matrix of normalized fit scores per cluster (**Figure 3A**). These scores could range from 1 to -1. Positive values indicate similar activity patterns, negative values indicate inverse activity patterns, and 0 indicates no relationship between activity patterns. Fit scores were bootstrapped from across neurons within a cluster to obtain a 95% confidence interval for fit scores. We further visualized the geometry of similar vs dissimilar event-related activity patterns via multi-dimensional scaling (**Figure 3B**).

**Figure 3.**
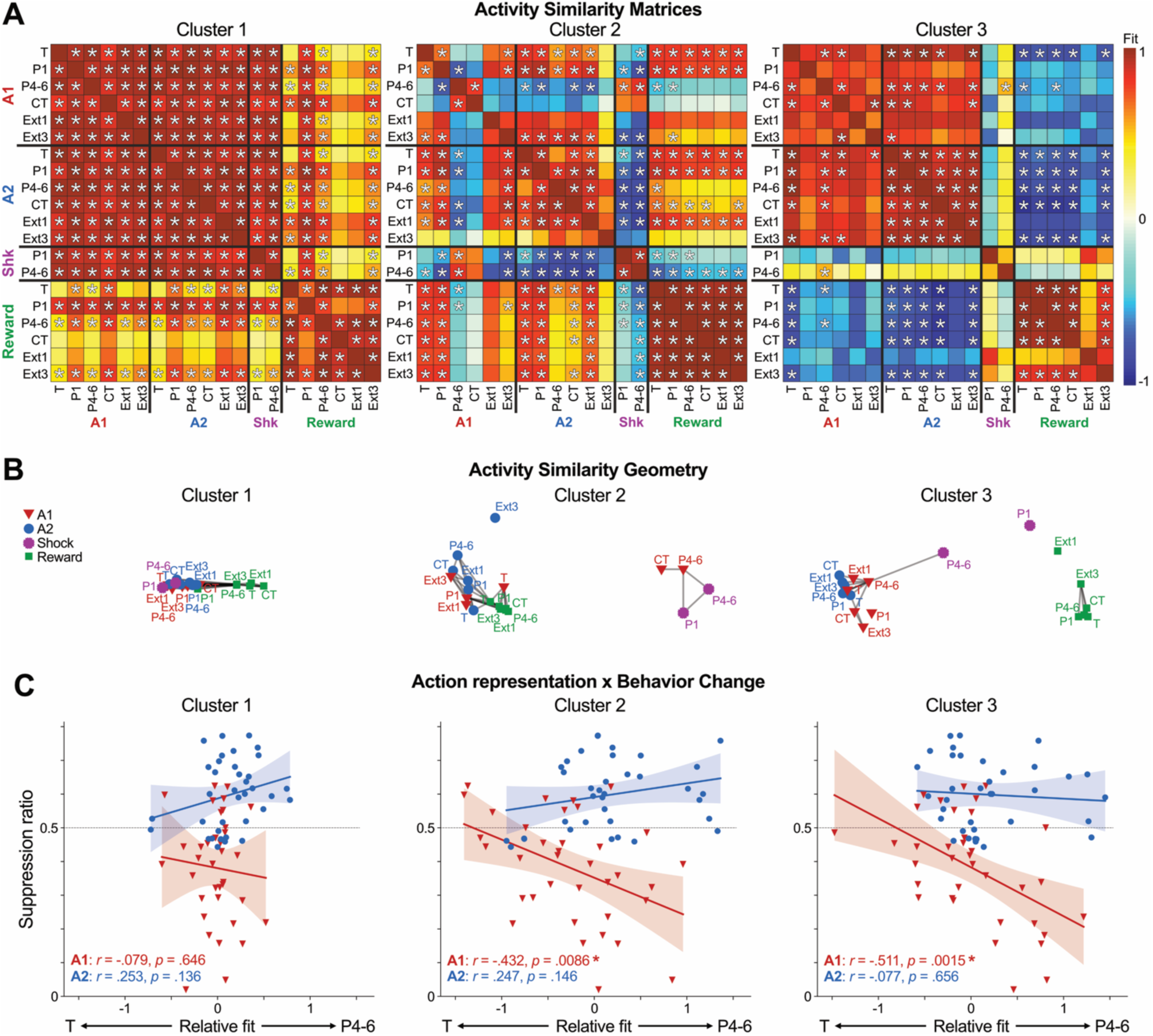
BLA clusters have dissociated representational geometries that track behavior change across punishment phases. **A** Similarity matrices of normalized activity patterns to actions (A1,A2) and outcomes (Reward, Shock [Shk]) across sessions (lever training [T], early punishment [P1], late punishment [P4-6], choice test [CT], Extinction 1, Extinction 3) per cluster. Fit values ranged from 1 (identical) to -1 (perfectly inverse). Asterisks (*) indicate where pairwise fits were significantly similar (>0) or dissimilar (<0), as determined via 95% bootstrapped fits. **B** Depiction of activity similarity across events and sessions in 2D space (multidimensional scaling of fit distance [1 - fit]) per cluster. Lines connecting nodes indicate significantly similar event-by-session activity patterns (as indicated in Figure 3A). **C** Relationships between changes in A1 and A2 coding (relative to late punishment [P4-6] vs pre-punishment [T]) and changes in lever-pressing relative to lever training (suppression ratio; no change = 0.5). Each dot represents relative fit for a mouse’s cluster activity and that mouse’s suppression for a given session (P1, P2, P3, CT, Ext1, Ext3). Shifts in Cluster 2 and Cluster 3 A1 coding towards/away from late punishment coding predicted A1 suppression in intervening and subsequent sessions. Shifts in A2 coding did not predict unpunished behavior in any cluster.

This analysis showed that not only do BLA ensembles differ in their activity patterns to task events (directions and magnitudes; **Figure 2**), but also in their coding principles (i.e., in the relationships between activity patterns across events and phases). Cluster 1 neurons operated in a constrained coding space, showing similar and stable phasic activity relative to the onset of actions and outcomes across the task. By contrast, Cluster 2 neurons exhibited stable, inverse activity patterns to aversive vs. appetitive outcomes, and a significant re-organization of activity patterns to punished A1 but not A2 across phases of the task. Pertinently, unpunished and pre-punished actions were initially similar to reward activity. As punishment was learned, punished actions became significantly dissimilar to unpunished actions and rewards, and instead became significantly similar to the shock punisher it led to. This reversed with punishment extinction. Finally, Cluster 3 neurons had relatively stable coding of actions (particularly unpunished A2) which were dissociated from its coding of rewards. Interestingly, shock punisher activity was not significantly similar to any other event except late punishment shock activity being similar to late punishment A1 activity.

These analyses indicate action-selective, contingency-dependent remapping of punished actions towards punishers in specific ensembles. However, these analyses do not directly show the relevance of this to individuals’ behavior across phases. To examine this, we assessed how recoding of actions per subject’s cluster related to behavior change across sessions (**Figure 3C**). This was conducted using self-normalized measures of activity change (relative similarity of a session’s action kernel to late vs pre-punishment per subject’s cluster) against a self-normalized lever-press ratio. To avoid circularity when assessing relative similarity of pre- and late punishment kernels to themselves, we only used relative fits for non-comparison sessions (Pun1-3, CT, Ext1-3). We found that the degree to which a subject avoided the punished action in a given session was strongly predicted by how much punished action activity had shifted towards late punishment activity patterns for Cluster 2 (*r*=-.432, *p*=.0086) and Cluster 3 neurons (*r*=-.511, *p*=.0015) but not Cluster 1 neurons (*r*=-.079, *p*=.646) (**Figure 3C**). There were no significant relationships between shifts in unpunished action coding and unpunished behavior for any cluster (|*r*|≤.253, *p*≥.136). This indicates that recoding of punished actions in BLA is strongly related to punishment-related behavior change across subjects.

Together, these findings indicate discrete, task-relevant patterns of activity in BLA neurons that predict behavior across phases of punishment learning, choice, and extinction. The disparate representational spaces of these clusters suggest non-equivalent coding principles across BLA neurons.

### Whole-population geometry of punishment and reward representations

A caveat of these analyses is that they do not show how BLA neurons collectively represent punishment learning because unclustered neurons were omitted from analyses and because findings were based on within-cluster analyses. Previous research has shown that BLA neurons, when analyzed collectively, encode broader functional dimensions (e.g., valence) beyond what individual BLA neurons individual encode (Kyriazi et al., 2018). We therefore examined how the collective population of BLA neurons, including unclustered neurons, represent punishment learning. To examine this, mean event-related activity from all 1,147 registered neurons across phases were used to extract the geometry of whole-population task-related activity (**Figure 4A**).

**Figure 4.**
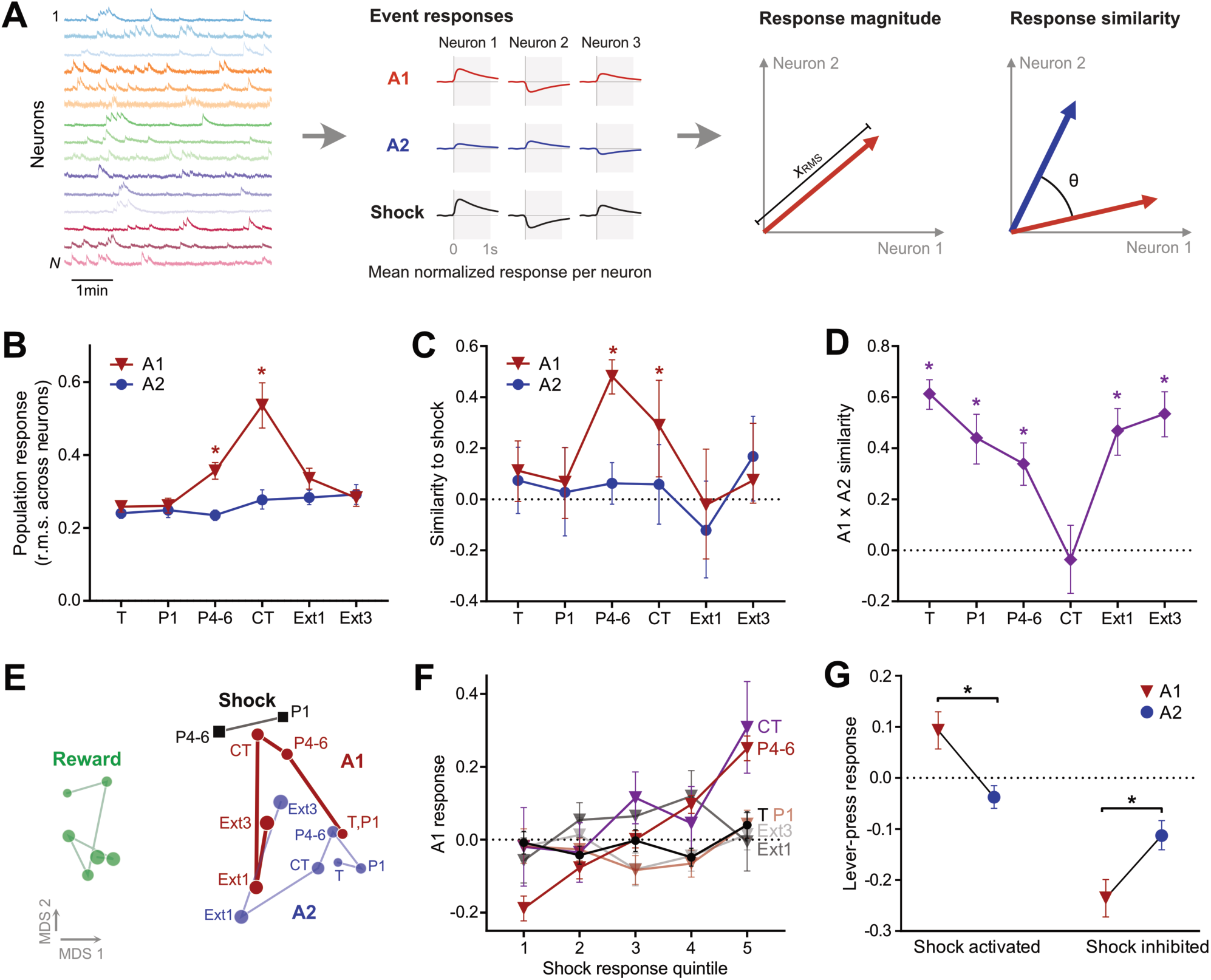
Punished-lever population responses rotate toward the shock representation across punishment learning. **A** Analysis schematic. Left: normalized activity traces from example BLA neurons in one session. Middle: for each neuron, the average trial-normalized response to each event (e.g., A1, A2, shock) from 0–1s after event onset (grey shading) was summarized as a single population response vector per event. Right: two measures derived from these vectors: response magnitude (the root mean square across neurons) and response similarity (cosine of the angle [θ] between a pair of population vectors). **B** Population response magnitude (root mean square across neurons) for A1 and A2 by session. Error bars = 95% bootstrapped CI across neurons. *p < 0.05, A1 vs A2. **C** Alignment of each event’s population response vector with the shock axis (Pun 4–6 shock response) by session. Error bars = 95% bootstrapped CI across neurons. *CI excludes zero. **D** Cosine similarity between the punished (A1) and unpunished (A2) press population vectors within each session (identical neurons within session). Error bars = 95% bootstrapped CI across neurons. *CI excludes zero. **E** Multidimensional scaling of all 20 session × event conditions (cosine distance, pairwise-complete neurons). The punished action (A1) traverses the space between unpunished actions and shock across sessions, while the unpunished lever press (A2) and reward remain relatively stationary. Axis units and rotation are arbitrary. **F** Punished action (A1) response by neuron quintile of Pun 4–6 shock response, separately for each session. Error bars = SEM across neurons. **G** Response of shock-activated and shock-inhibited neurons to punished (A1) and unpunished (A2) levers in Pun 4–6; neurons split by the sign of their Pun 4–6 shock response (n = 310 activated, 254 inhibited). Error bars = 95% bootstrapped CI across neurons. *p < 0.05, paired test between actions. Sessions: T = lever-press training; P1 = punishment day 1; P4–6 = punishment days 4–6 (pooled); CT = choice test; Ext1 and Ext3 = extinction days 1 and 3.

Punishment increased the magnitude of population activity associated with punished A1 (**Figure 4B**). Magnitude increased from lever-press training to Pun 4–6 (95% CI = 0.052 to 0.119, *n* = 269), increased further from Pun 4–6 to choice test (95% CI = 0.115 to 0.264, *n* = 163), and then decreased from choice test to Ext 3 (95% CI = −0.343 to −0.133, *n* = 95) (all q < 0.002), by which point it no longer differed from training (95% CI = −0.021 to 0.053, p = 0.39, *n* = 140). Indeed, the increase for the punished press exceeded that for the unpunished A2 press at both Pun 4–6 (95% CI = 0.061 to 0.143, *n* = 269) and choice test (+0.254, 95% CI = 0.169 to 0.339, *n* = 145). As the choice test and extinction sessions contained no shock at all, the increase at choice and reversal at extinction cannot be due to the presence of the punisher.

If BLA comes to represent an action partly in terms of its outcome, then the A1 press population vector should become increasingly aligned with the shock vector as the A1–Shock contingency is learned. To assess this, we projected each condition onto a fixed shock axis (**Figure 4C**). A1 alignment with the Pun 4–6 shock vector did not differ from chance during lever-press training (95% CI = −0.009 to 0.228, *p* = 0.22, *q* = 0.32, *n* = 265) or Pun 1 (95% CI = −0.075 to 0.203, p = 0.81, q = 0.87, n = 194) but emerged by Pun 4–6 (95% CI = 0.412 to 0.546, p = 0.0002, q = 0.001, n = 564), remained elevated at choice test (95% CI = 0.088 to 0.466, p = 0.0004, q = 0.001, n = 162), and returned to chance during extinction (Ext 1, 95% CI = −0.234 to 0.196, p = 0.83, q = 0.88, n = 129; Ext 3, 95% CI = −0.015 to 0.298, p = 0.083, q = 0.15, n = 166). The same pattern was observed with the independently measured Pun 1 shock vector: alignment of the punished press rose from lever-press training (0.17, q = 0.11, n = 224) to Pun 4-6 (0.29, q = 0.001, n = 195) and fell to zero by Ext 3 (−0.10, q = 0.44, n = 103), with both the increase (95% CI = 0.07 to 0.40, n = 154) and decrease (95% CI = 0.16 to 0.60, n = 88) present in neuron-paired testing. The increase at Pun 4–6 was present in neuron-paired analyses restricted to cells recorded at both time points (95% CI = 0.06 to 0.40, p = 0.006, q = 0.017, n = 154), as was the subsequent decline during extinction (95% CI = −0.61 to −0.16, p = 0.0008, q = 0.003, n = 88). This alignment was specific to the punished action. At Pun 4–6, the punished press was significantly more shock-aligned than the unpunished press measured in the same neurons (difference = 0.42, 95% CI = 0.32 to 0.51, p = 0.0004, q = 0.001, n = 564; **Figure 4C**).

We then asked how this learning altered the relationship between representations of the two actions (**Figure 4D**). A1 and A2 press vectors were similar during lever-press training (cosine = 0.61, 95% CI = 0.56 to 0.67, n = 508), progressively less so across punishment (Pun 1, 0.44, 95% CI = 0.34 to 0.53, n = 315; Pun 4–6, 0.34, 95% CI = 0.25 to 0.42, n = 573), orthogonal at the choice test (−0.04, 95% CI = −0.17 to 0.10, p = 0.53, q = 0.64, n = 267), and reconverged during extinction (Ext 1, 0.47, 95% CI = 0.37 to 0.56, n = 291; Ext 3, 0.53, 95% CI = 0.44 to 0.62, n = 269); all except the choice test differed from zero (p < 0.0002, q < 0.001). Both the separation at choice test relative to training (95% CI = −0.85 to −0.45, q = 0.001, n = 145) and the subsequent reconvergence by Ext 3 (95% CI = 0.36 to 0.94, q = 0.003, n = 95) survived neuron-paired analyses. The same separation and reconvergence was observed in the embedded population state space (**Figure 4E**). Thus, BLA representations of the two actions progressively diverged with punishment learning, were maximally distinct when animals chose between them, and reconverged when the punishment contingency was extinguished.

Our peri-event analyses indicated considerable variation in individual BLA neuron responses during punishment learning, including excitatory versus inhibitory responses to shocks and actions. To determine which neurons drove alignment between the punished-action and shock population activities, we ranked neurons by their shock response and examined their punished-press responses (**Figure 4F**). At Pun 4–6, punished-press responses increased monotonically across shock-response quintiles. Crucially, the same relationship held when computed across all neurons without binning: a neuron’s A1 response was strongly correlated with its shock response (r = 0.52, 95% CI = 0.46 to 0.58, p = 0.0002, n = 564). This correlation grew with punishment and reversed with extinction; it was weaker during lever-press training (r = 0.20; difference = 0.39, 95% CI = 0.26 to 0.52, p = 0.0004, n = 265) and had returned to that level by Ext 3 (r = 0.20; difference from Pun 4–6 = −0.37, 95% CI = −0.52 to −0.21, p = 0.0004, n = 166). It was also specific to the punished action. At Pun 4–6 the A1 response correlated with shock response significantly more strongly than the unpunished press did (0.52 versus 0.13; difference = 0.40, 95% CI = 0.30 to 0.50, p = 0.0004, n = 564), whereas the two actions did not differ at other stages, including choice test (difference = 0.20, 95% CI = −0.07 to 0.45, p = 0.16, n = 162). The correlation reflected matching response polarity. At Pun 4–6, shock-activated neurons were generally activated by the punished press (95% CI = 0.06 to 0.13, n = 310) and shock-inhibited neurons were generally inhibited by it (95% CI = −0.27 to −0.20, n = 254; **Figure 4G**). This was not observed for unpunished A2 presses, which evoked inhibition in both shock-activated (95% CI = −0.06 to −0.02) and shock-inhibited neurons (95% CI = −0.14 to −0.08). These differences between A1 and A2 were significant (shock-activated: 95% CI = 0.097 – 0.165, p = 0.0004, n = 310; shock-inhibited: 95% CI = -0.163 – -0.081, p = 0.0004)

Further analyses tested non-specific accounts of this correlation. A gain, salience, or excitability account of this action–punisher correlation predicts that is due to matched response magnitudes, not matched signs. However, the action–punisher correlation was weaker on absolute than on signed responses (r = 0.20 versus 0.52). Furthermore, it was not related to neuronal responses to reward as factoring in each neuron’s reward response left the correlation unchanged (partial r = 0.53 versus r = 0.52). Finally, it was not due to neurons with extreme shock responses, because it survived rank transformation (Spearman ρ = 0.52) and removal of the most extreme 20% of shock responders (r = 0.38).

Taken together, these results show that punishment learning progressively reorganizes the representation of a punished action across BLA neurons. The punished action separates from the unpunished action and becomes aligned with shock population activity. This alignment reflects the signed structure of the shock population activity: neurons activated by shock become increasingly activated by the punished action, whereas neurons inhibited by shock become increasingly inhibited by the punished action. This organization emerges across learning and reverses when the action–shock contingency is extinguished. Thus, the BLA representation of a punished action is progressively transformed toward the representation of its aversive outcome.

A critical question is what this transformation represents. One possibility is that the punished action simply acquires negative value. Reinforcement-learning accounts commonly formalize punishment as a negative outcome that reduces action value (Sutton and Barto, 2018). The population code is then read out as movement along a single good-to-bad axis. A scalar value code predicts that reward and shock occupy opposite poles of a common axis. They did not. Shock and reward occupied nearly orthogonal rather than opposite population directions (cosine = 0.09 [85°], 95% CI = 0.006 to 0.176, n = 564; **Figure 5A**). Moreover, even if opponent coding existed in the BLA, our data constrain its magnitude during punishment. The displacement of the punished-press vector from lever-press training to Pun 4–6 had a large positive shock component (cosine = 0.494, 95% CI = 0.385 to 0.592, n = 265) but minimal reward component (cosine = −0.018, 95% CI = −0.165 to 0.125) (**Figure 5B**). Expressed differently, from the confidence intervals, punishment learning plausibly subtracted none and at most a third as much appetitive component as it added aversive.

**Figure 5.**
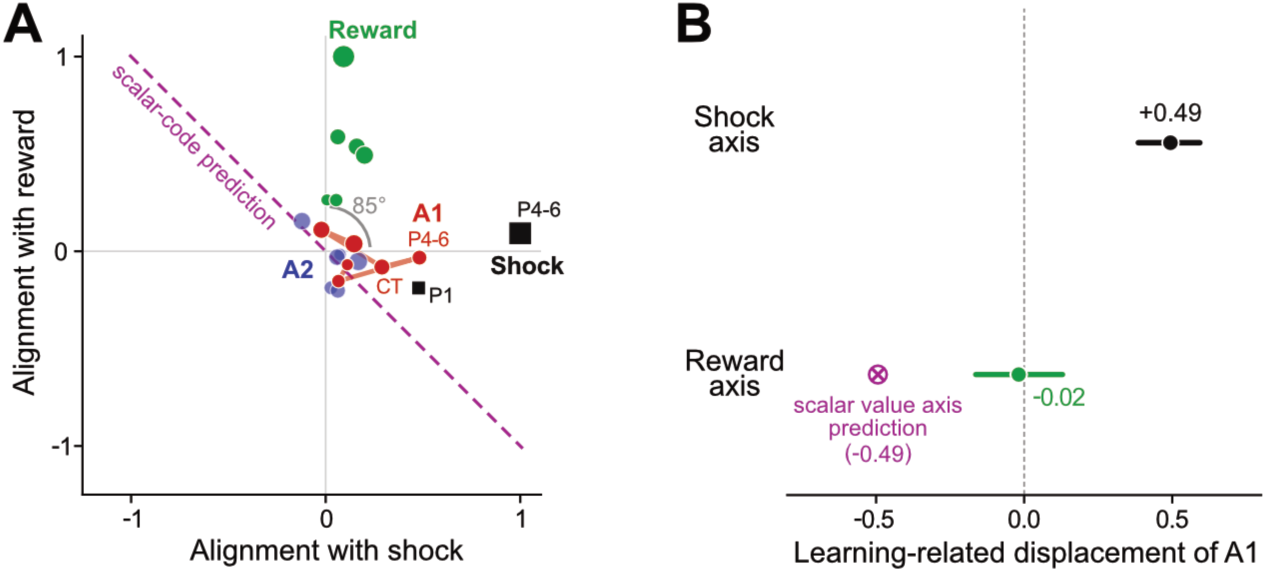
The punished lever moves along the shock axis without an opposing shift along the reward axis. **A** Every session × event condition placed in the plane defined by the shock axis and the reward axis (population response vectors to shock vs. reward consumption at Pun 4–6), plotted as the cosine similarity of its own response vector with each axis (pairwise-complete neurons). Marker size increases with session order; the red line traces the A1 trajectory across sessions. Grey arc indicates the near-orthogonal 85° angle between reward and shock axes. Magenta dashed line indicates the displacement axis predicted by a scalar value code. **B** Learning-induced displacement of the punished action (A1) population vector at Pun 4–6, defined as the change from pre-punishment (Pun 4–6 minus T for n = 265 neurons imaged in both phases) along the shock axis (black, +0.49) and reward axis (green, −0.02). Error bars = 95% bootstrapped CI across neurons. Purple circle with cross indicates predicted displacement along reward axis if punished action (A1) displacement occurred according to a scalar value code (given shock axis displacement).

## Discussion

Here we used longitudinal in vivo calcium imaging to study BLA neuronal activity during punishment learning. Consistent with previous descriptions of valence-selective populations in BLA (Namburi et al., 2015; Beyeler et al., 2016; Kim et al., 2016; Piantadosi et al., 2024), neurons fell into profiles with dissociated responses to reward and punishment, and some of these neurons acquired a unique response to the punished action across learning. Population activity associated with the punished action increased in magnitude, progressively separated from that associated with the unpunished action, and became aligned with the representation of the punisher. This transformation developed across training, was selective for the punished action, persisted when animals chose between the actions in the absence of punishment, and reversed when the action–punisher contingency was extinguished. Moreover, this alignment reflected the signed population structure of punisher representation: neurons activated by shock tended to be activated by the punished action, whereas neurons inhibited by shock tended to be inhibited by it. Thus, punishment learning progressively shifted the population representation of the punished action toward that of the punisher itself.

This transformation has a clear precedent in associative learning theory. Learning establishes relationships between cues or actions and representations of their outcomes. In Pavlovian conditioning this relation is between stimulus and outcome, whereas in instrumental learning it is between response and outcome (Dickinson and Mackinosh, 1978; Mackintosh, 1983). Consistent with this idea, Pavlovian fear conditioning transforms BLA representations of conditioned stimuli toward those of the unconditioned stimuli they predict (Grewe et al., 2017; Zhang and Li, 2018). Our results extend this principle from cues to actions. Interestingly, unlike the rapid reorganization of BLA choice representations produced by changes in punishment risk (Piantadosi et al., 2026), this transformation emerged across learning, persisted in the absence of punishment, and reversed during extinction. Importantly, it preserved the signed structure of the punisher representation: neurons excited by shock became excited by the punished action, whereas neurons inhibited by shock became more inhibited by it. Associative learning preserved the internal structure of that representation, including the direction of individual neuronal responses. Thus, although neuronal excitability may influence memory allocation (Josselyn and Frankland, 2018), preferential recruitment alone cannot explain this correspondence between action and outcome representations.

The choice test was particularly informative because the punished action continued to recruit the punisher representation when punishment was no longer present. Indeed, the most reliable population response to the punished action occurred when animals chose between the two actions in the absence of shock. Thus, the transformation cannot be explained simply by the immediate sensory consequences of punishment. Instead, a learned representation of the aversive outcome was expressed when that outcome was absent but relevant to decision-making. This accords with evidence that BLA encodes outcome-specific incentive information to guide instrumental choice (Balleine and Killcross, 2006; Balleine, 2019). In appetitive instrumental learning, Parkes and Balleine (Parkes and Balleine, 2013) further distinguished the encoding of incentive value in BLA from its retrieval during choice, which depended on insular cortex. Our results complement this distinction by showing that, during choice, BLA population activity associated with the punished action nevertheless contains a representation of its learned aversive outcome.

Punishment also provides an informative assessment of how multiple outcomes of the same action are represented, because the action retains its association with reward while acquiring an aversive outcome. Balleine and Killcross (2006) proposed that BLA represents the incentive properties of specific motivationally significant outcomes rather than general affective value. Consistent with this proposal, BLA activity associated with instrumental actions contains information about prospective rewarding outcomes, including their identity, value, and expectancy (Courtin et al., 2022; Riffault et al., 2024). Reinforcement-learning models provide a different representational possibility: appetitive and aversive outcomes can contribute with opposite signs to a scalar action value. If BLA instantiated a common value representation, punishment learning should move an action representation towards shock and commensurately away from reward. We observed the former but not the latter. The punished action acquired a strong shock component but did not lose its reward component, while reward and shock themselves occupied largely orthogonal rather than opposing population directions. So, punishment added a new aversive outcome to the action representation rather than simply subtracting value from an existing representation. In associative terms, punishment did not simply alter a scalar property attached to the response; it altered the represented outcomes of that response. The same action could retain a representation of reward while acquiring a separable representation of the punisher. Such an architecture preserves specific information about each outcome, leaving any integration for choice to downstream regions, potentially including ventral striatum (McNally, 2021; Piantadosi et al., 2026) and/or insular cortex (Parkes and Balleine, 2013).

This architecture also fits a well-established property of punishment learning: punishment does not erase the reward learning that originally supported an action. Punished responding can renew when animals are tested outside the punishment context, and contextual control can be specific to the punished response (Marchant et al., 2013; Bouton and Schepers, 2015). These findings suggest that, like extinction, punishment establishes additional learning that modifies action selection while leaving earlier reward learning intact (Khoo et al., 2017; Bouton et al., 2021). Our results reveal a corresponding organization in BLA. Reward- and punisher-related information about the same action remained separable: the aversive component tracked the prevailing action– punisher contingency, whereas the appetitive component persisted through both its acquisition and extinction. Thus, the population code preserves distinct consequences of the same action rather than replacing its appetitive representation with an aversive one. Such an organization could allow context, motivational state, and other variables to regulate action selection without requiring the underlying action–outcome representations themselves to be rewritten.

### Methodological considerations

First, population geometry identifies relationships among BLA representations but does not establish their causal role in behavior. The causal role of amygdala in human and rodent punishment behavior is well established (Killcross et al., 1997; Jean-Richard-Dit-Bressel and McNally, 2015; Orsini et al., 2015; Truckenbrod et al., 2023; Gregory et al., 2026; Piantadosi et al., 2026), but our findings do not show that the population transformations detected here cause suppression of the punished action or how downstream structures read out the distinct reward and shock dimensions to control action selection. Relatedly, the peri-action population activity may contain contributions from action execution, sensory feedback, and outcome retrieval. The selective emergence of punisher alignment for the punished action, including during choice in the absence of shock, strongly implicates learned outcome information, but higher temporal resolution or causal manipulation of the relevant population activity will be required to determine when and how this outcome information contributes to action selection. Second, GCaMP7f expression was driven by the hSyn promoter rather than being restricted to a genetically defined neuronal population. We used hSyn deliberately to sample broadly across BLA neurons but cannot identify the contributions of specific cell types. Third, food pellets and shock constitute a single pair of appetitive and aversive outcomes. Their near-orthogonal organization argues against a single appetitive-to-aversive BLA axis in this task but does not imply that all rewarding and aversive outcomes occupy independent dimensions. Fourth, longitudinal registration yielded overlapping rather than identical neuronal populations across the experiment. Indeed, only a small number of neurons were detected in every session. The critical effects survived analyses restricted to neurons recorded across the relevant stages, but larger populations tracked across the entire experiment would permit stronger tests of how individual neurons contribute to these population transformations. Finally, we used only male mice. Sex differences have been observed in punishment learning (Orsini and Setlow, 2017; Chowdhury et al., 2019; Orsini et al., 2022), so assessing the population geometry described here in females will be important.

## Conclusions

Together, these findings identify a representational architecture in which punishment changes what an action is expected to cause without simply reversing its value. Through punishment learning, an initially rewarded action acquires an additional representation of its aversive outcome while retaining its representation of reward. More generally, BLA population activity can preserve opposing consequences of the same action rather than collapsing them onto a common value dimension. Such an architecture allows the same action to remain associated with reward and punishment, while context, current goals, and downstream circuits determine which consequence controls choice.

## Acknowledgments

This work was supported by Australian Research Council Discovery Project grants DP220100040 and DP250100345 to GPM, and Australian Research Council Future Fellowship FT260101559 to PJRDB.

## Author contributions

Conceptualization: GPM, PJRDB, SK; Methodology: NH, EC, EZM, COHB PJRDB, GPM; Investigation: NH, EAC; Visualization: PJRDB, GPM; Funding acquisition: GPM; Project administration: GPM; Supervision: GPM, SK, PJRDB. Writing – original draft: GPM, PJRDB; Writing – review & editing: All authors.

